# Pinpointing the distinctive impacts of ten cover crop species on the resident and active fractions of the soil microbiome

**DOI:** 10.1101/2022.06.10.495641

**Authors:** Sara Giulia Cazzaniga, Lize Braat, Sven van den Elsen, Carin Lombaers, Johnny Visser, Jose G. Maciá-Vicente, Joeke Postma, Liesje Mommer, Johannes Helder

## Abstract

Cover crops are used in agriculture to minimise soil erosion, prevent nutrient leaching and increase soil organic matter content. Cover crops can also be grown to stimulate the soil microbial community to improve soil biological conditions. Despite their widespread use, little is known about the impact of different cover crop species on the composition and activity of the soil microbiome. Here we investigate the effect of distinct cover crop species on the rhizosphere microbiome and to characterise both the resident (DNA-based) and the active (RNA-based) fractions of the bacterial, fungal, protist and metazoan communities in the cover crops rhizosphere. We conducted a field experiment using 70-litre bottomless containers in which we grew ten monocultures of commonly used cover crop species belonging to five plant families, and an unplanted control treatment (fallow). The total DNA and RNA were extracted from soil and the bacterial, fungal, protistan and metazoan communities were characterized using Illumina MiSeq sequencing. We found that all cover crop species significantly impacted the resident and active microbial community composition. Moreover, cover crops showed distinct selection strengths on their rhizospheres. Together with borage (Boraginaceae), oilseed radish (Brassicaceae) was shown to provoke the strongest microbial shifts, in part attributable to a promotion of the bacterial family Pseudomonadaceae and a repression of Microascaceae in the rhizosphere. Lentil (Fabaceae) showed enrichment in fungal taxa, including Trichocomaceae and fungal members of the Glomerales order, whereas black oat, hybrid ryegrass (both Poaceae) and marigold (Asteraceae) induced relatively mild changes in the soil microbial communities. Predominantly, differences in selection strengths were consistent among the four organismal groups under investigation and were observed in both the active and resident communities. Our work provides a broad baseline for the effects of cover crops on four organismal groups, which may facilitate future cover crop selection to advance soil health.

## 1. INTRODUCTION

From the 1960s onwards, agricultural intensification has led to higher and more stable crop yields with a fraction of the labour inputs previously needed (Normile and Mann, 1999). However, intensive agriculture also carries negative effects on soil health, including degradation of the physical, chemical and biological properties of soils (Tsiafouli et al., 2015; Banerjee et al., 2019). A recent report states that 60-70% of the soils within the European Union are classified as unhealthy as a result of current agricultural practices (Veerman et al., 2020). Reconsideration and adjustments of these practices are needed to reverse this undesirable phenomenon.

A number of soil management practices have been shown to be effective in improving soil health while maintaining acceptable crop production levels (Vukicevich et al., 2016; Schrama et al., 2018; Eyhorn et al., 2019). Among these, cover cropping - the cultivation of fast-growing non-economic plants between the harvest of the main crop and the sowing of the next main - is implemented to minimize nutrient leaching and soil erosion, and to increase the soil organic matter content (Blanco-Canqui et al., 2015; Kaye and Quemada, 2017; Wick et al., 2017). Cover cropping also has a positive effect on the biological condition of soils as it can boost the activity and abundance of the soil microbes (Vukicevich et al., 2016; Wick et al., 2017; Kim et al., 2020). Potential downsides of cover crops include their potential function as reservoirs for pests and pathogens that facilitates them to bridge a non-favourable period in their life cycle (Bakker et al., 2016; Walder et al., 2017), and their action as weeds in the next cropping season (Wayman et al., 2015). However, as long as these risks are mitigated, many plant species can be considered for cover cropping.

Cover crops steer the soil microbiome via the release of root exudates during their growth, and by residue decomposition after the termination of the crop. In this study, we focus on the influence of growing cover crops on the soil microbiome during plant growth. It is estimated that up to 40% of the carbon fixed by plants is released into the rhizosphere through root exudates which can modulate the microbial community directly associated with roots (Berendsen et al., 2012). The steering of the local microbiome by plants is largely dictated by the composition of these exudates (Berg and Smalla, 2009), which, in turn, is greatly determined by plant phylogeny at the plant family level but in many cases also at species-specific and even -subspecific levels (Pascale et al., 2020). Root exudates include both primary metabolites, such as fatty and aromatic organic acids with proven effects on root-associated bacterial communities (Zhalnina et al., 2018), and secondary metabolites, such as triterpenes, coumarins and benzoxazinoids, which have also been shown to affect the root microbiome (Stringlis et al., 2018; Cotton et al., 2019; Huang et al., 2019).

Although rhizosphere communities have been characterized for a substantial number of plants, and, in particular, economic crops (Fitzpatrick et al., 2018; Yadav et al., 2018), our knowledge of the microbial signatures of cover crop species is rather crude. For example, phospholipid fatty acid (PLFA) profiles across eight cover crop species suggested an increased abundance of arbuscular mycorrhiza (AM) in oat and cereal rye (Poaceae), whereas non-AM fungi were stimulated by hairy vetch (Fabaceae) (Finney et al., 2017). Bacq-Labreuil et al. (2019) also used PLFA profiles to investigate the impact of eight growing cover crops on the microbial community of bulk soil, and they found that phacelia (Boraginaceae) stimulated saprophytic fungi. In a more detailed study using a next-generation sequencing approach focused on the impact of a single cover crop species - oilseed rape (*Brassica napus*) - on the resident and the active fractions of the bacterial and fungal communities in the rhizosphere and in the plant roots (Gkarmiri et al., 2017). Proteobacteria were highly active in the oilseed rape rhizosphere, whereas Actinobacteria, despite of their abundance, had a low activity. These studies illustrate the potential of cover crop species to influence the soil microbiome. However, the available knowledge is mostly generated at a high taxonomic level, limited to few cover crop species, and seldomly focused on the rhizosphere microbiome.

In addition to plant identity, bacterial and fungal soil communities are co-shaped by the activity of major consumers of these primary decomposers, *i*.*e*., protists and metazoans (Gao et al., 2019; Mielke et al., 2022). In the temperate agricultural systems, primary consumers-biomass in the top layer of arable fields is typically 40 to 100 times smaller than the bacterial and fungal biomass (Pausch et al., 2018). However, several studies demonstrated the high metabolic activity and growth potential of these primary consumers (Foissner, 1999; Finlay et al., 2000; Liu et al., 2017). Therefore, major representatives of the primary consumers should be taken along to achieve a proper understanding of the shaping of the rhizobiome.

Recent studies underlined the relevance of including the transcriptionally active fraction of the microbiome as a relevant parameter to assess the functioning of soil microbial communities (Ofek et al., 2014; Harkes et al., 2019). A substantial part of the soil microbial community is known to be dormant (Fierer, 2017). This ‘microbial seed bank’ as it was referred to by Lennon and Jones (2011) may comprise up to 80% of the cells and about 50% of the taxa in bulk soils. Hence, it is essential that - next to the resident community - also the active fractions in the rhizobiome of cover crops should be investigated.

Here, we present a field experiment in which we characterized the impact of ten cover crops belonging to five different plant families on both the resident and the active fractions of the bacterial, fungal, protist and metazoan rhizosphere communities. We investigate whether distinct cover crop species have a differential impact on the resident and the active fractions of the primary decomposers (bacteria and fungi) in the rhizosphere, as well as of major representatives of their consumers (protists and metazoa). This overall question was addressed both quantitatively - do the selection strengths of cover crops on the soil microbiome differ? - and qualitatively - do cover crops differ in the microbial taxa that they promote and/or repress? A better understanding of the specific microbial signatures of several cover crops would allow for applications beyond the current, general, scope. It would facilitate the selection of particular cover crops to steer the soil microbial community in such a way that they may contribute to restoring healthy soil.

## 2. MATERIALS AND METHODS

### 2.1 Experimental design

A field experiment was carried out at the Wageningen University and Research experimental farm ‘Vredepeel’, located in the southeast of the Netherlands. The experiment consisted of 11 treatments, including ten cover crop species and an unplanted control (fallow), each replicated eight times. Cover crop treatments included widely used cover crop cultivars, as well as a prospective oilseed radish cultivar referred to as E1039. Bottomless containers (70L) were randomly positioned in eight blocks (a total of 88 plots). The containers were dug into the field in such a way that there was no height difference between the soil surface in the buckets and the surrounding soil. The containers were filled with topsoil (20 cm) originating from subplots of a nearby long-running field experiment ‘Soil Health Experiment’ (SHE) (Korthals et al., 2014). Each block of containers was filled with the topsoil originating from one of the SHE subplots. Half of the SHE subplots were managed following organic practices and the other half following conventional practices until 2017. It is noted that all subplots received the same soil management in the two years before this field experiment (a single application of cattle slurry per year) (field management data provided in Suppl. Table 1). In each block, ten containers were sown with single cover crop cultivars (Table 1), and one container was kept fallow. Cover crop seeds were sown at the end of July 2019 (sowing densities shown in Table 1), and weeds were removed manually during the duration of the experiment. In 14 containers cover crop growth was negatively affected by drought in the late summer of 2019 (weather data provided in Suppl. Table 2) and excluded from the experiment. Thus, 74 of the original 88 plots were sampled at the end of the experiment (list in Suppl. Table 3).

**Table 1.**
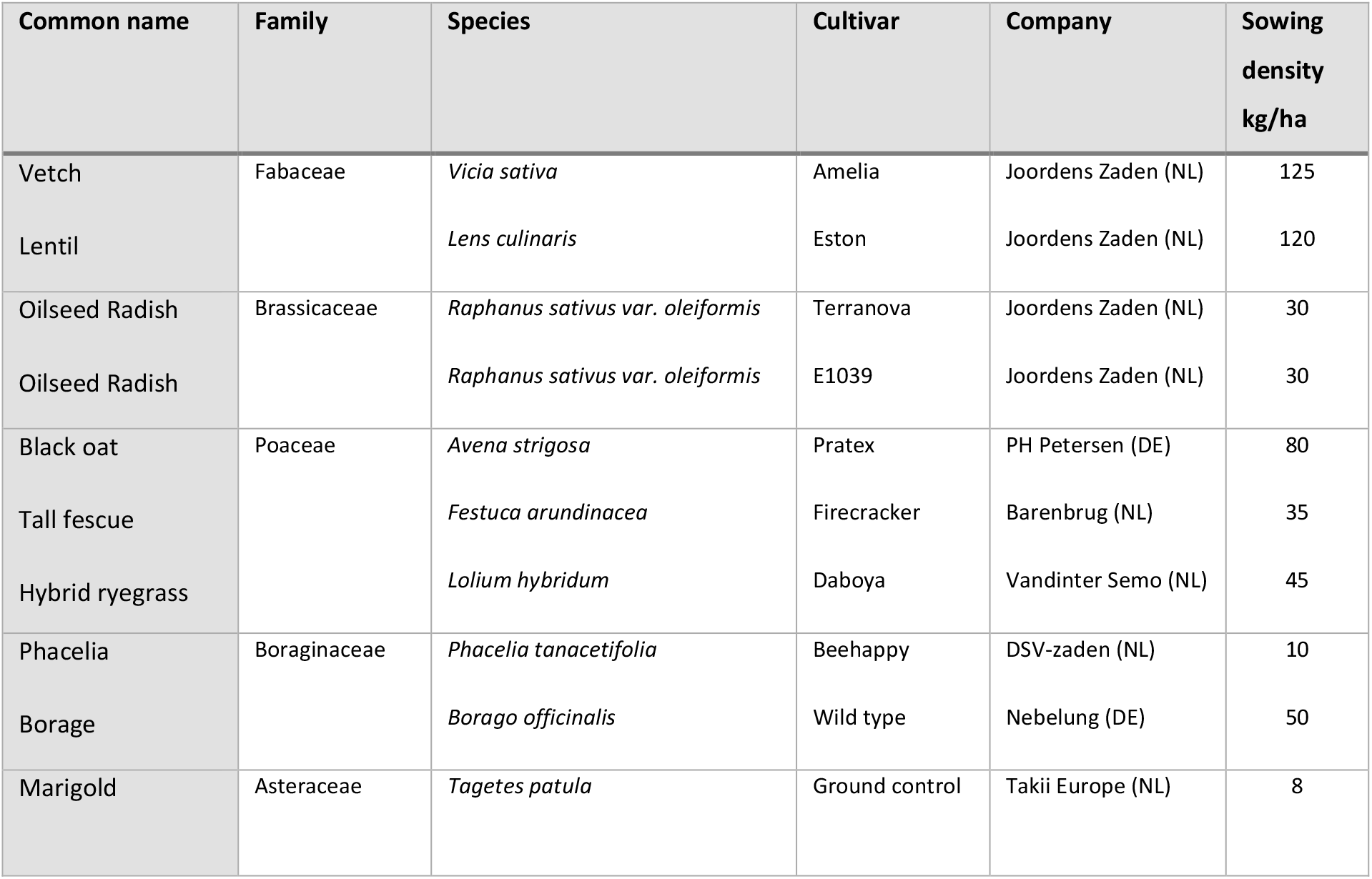
Details of the cover crop species used in this study, including taxonomic affiliation, the origin of seeds, and sowing density.

| Common name | Family | Species | Cultivar | Company | Sowing density kg/ha |
| --- | --- | --- | --- | --- | --- |
| Vetch | Fabaceae | *Vicia sativa* | Amelia | Joordens Zaden (NL) | 125 |
| Lentil |  | *Lens culinaris* | Eston | Joordens Zaden (NL) | 120 |
| Oilseed Radish | Brassicaceae | *Raphanus sativus var. oleiformis* | Terranova | Joordens Zaden (NL) | 30 |
| Oilseed Radish |  | *Raphanus sativus var. oleiformis* | E1039 | Joordens Zaden (NL) | 30 |
| Black oat | Poaceae | *Avena strigosa* | Pratex | PH Petersen (DE) | 80 |
| Tall fescue |  | *Festuca arundinacea* | Firecracker | Barenbrug (NL) | 35 |
| Hybrid ryegrass |  | *Lolium hybridum* | Daboya | Vandinter Semo (NL) | 45 |
| Phacelia | Boraginaceae | *Phacelia tanacetifolia* | Beehappy | DSV-zaden (NL) | 10 |
| Borage |  | *Borago officinalis* | Wild type | Nebelung (DE) | 50 |
| Marigold | Asteraceae | *Tagetes patula* | Ground control | Takii Europe (NL) | 8 |

**Table 2.** PERMANOVA analysis with Bray-Curtis dissimilarity metric to assess the variation explained by the cover crop treatments on the resident (DNA) and active (cDNA) communities separately, of each of the four organismal groups.

| ORGANISMAL GROUP | RESIDENT (DNA) |  |  | ACTIVE (cDNA) |  |  |
| --- | --- | --- | --- | --- | --- | --- |
| | $R^2$ | n | p-value | $R^2$ | N | p-value |
| bacteria | 0.40 | 73 (-1) | 0.001 | 0.40 | 74 | 0.001 |
| fungi | 0.44 | 74 | 0.001 | 0.51 | 70 (-4) | 0.001 |
| protists | 0.37 | 73 (-1) | 0.001 | 0.37 | 74 | 0.001 |
| metazoa | 0.33 | 68 (-6) | 0.001 | ND* | ND* | ND* |

$R^2$ = fraction of the variation explained by the experimental factor cover crop, n = the number of samples included for the test. Between brackets, the number of samples removed due to low sequencing coverage is given. Dissimilarity is significant for p-values < 0.01. ND = not determined

### 2.2 Soil sampling

Soil samples were collected in October 2019, approximately two and a half months after the sowing of cover crops. At the harvest, 4-10 plants, depending on the plant size, were randomly collected from each container, uprooted and shaken to discard non-rhizosphere soil. Plant samples were transported to the nearby laboratory, where rhizosphere soil was collected by brushing off the soil adhering to the roots. Fallow soil from the control container was collected with an auger (15 mm Ø x 20 cm depth). Three cores were sampled for each fallow container, and after thorough mixing and sieving (mesh size 5 mm), a subsample of 10 g was collected. Rhizosphere and fallow soil samples were transferred to clean Ziplock plastic bags, snap-frozen in liquid nitrogen, kept on dry ice during transport and subsequently stored at −80 °C at the Laboratory of Nematology.

### 2.3 Nucleic acids extraction and sequencing

Soil total DNA and RNA were extracted simultaneously following a protocol optimised for 2 g soil (Harkes et al., 2019). This extraction method comprises bead beating, precipitation of humic acids with an ammonium aluminium sulphate solution, and phenol-chloroform extraction. cDNA was synthesised from the extracted RNA using a Maxima First Strand cDNA Synthesis Kit for RT-PCR (Fermentas, Thermo Fisher Scientific Inc., USA) following the manufacturer’s instructions. In preparation for the first step of the library construction, all DNA and cDNA samples were diluted to 1 ng μl^-1^ and 0.1 ng μl^-1^, respectively. Following (Harkes et al., 2019), the library was generated in a two-step PCR procedure. The first step consisted of the amplification of organismal group-specific 16S and 18S rRNA regions. To this end, locus-specific primers extended with an Illumina read area and an appropriate adapter were employed that targeted the V4 region of 16S of bacteria, and the V9, V7-V8, V5-V7 of 18S of protozoa, fungi and metazoa, respectively (Suppl. Table 4). For PCR amplification, 3 μl of the diluted samples were used as templates. PCR was carried out with the following temperature profile: 3 s at 95 °C; followed by 39 cycles of 10 s at 95 °C, 20 s at 55 °C, and 20 s at 72 °C; and a final extension step of 5 s at 72 °C. All reactions in the first PCR step were done in triplicate, and PCR products were pooled per sample and organismal groups. The second PCR step was performed using 40x dilutions of the amplicons from the first PCR as templates. The second PCR was used to attach the sample-specific Illumina index combination, used for multiplexing the samples upon pooling, and the Illumina sequencing adapter to the amplicons of the first PCR. The following temperature profile was employed: 3 s at 95°C; followed by 10 cycles of 10 s at 95°C, 30 s at 60°C, and 30 s at 72°C; and a final extension step of 5 s at 72°C. A random selection of products of the first and second PCR steps was checked on an agarose electrophoresis gel to ensure that the amplifications were successful in producing amplicons of the expected size. Lastly, PCR products were pooled together and sent out for sequencing to Useq (Utrecht Sequencing Facility, Utrecht, The Netherlands). Illumina MiSeq sequencing was performed using a 2 × 300 bp V3 kit.

### 2.4 Pre-processing of the sequencing data

Sequencing data were demultiplexed and subset into the four organismal groups based on their locus-specific primer sequences, using a custom Python script. Sequencing reads were pre-processed in QIIME2 (Bolyen et al., 2019). Read picking was carried out with the QIIME2 DADA2 denoising algorithm (Callahan et al., 2016) separately for each organismal group. For bacteria, protists and fungi, paired-end reads were merged and used to generate amplicon sequence variants (ASVs) (Callahan et al., 2017). For metazoa, only the forward reads were used to generate ASVs.

Taxonomic assignment of ASVs was carried out using the q2-feature-classifier plugin and *classify-sklearn* function (Pedregosa et al., 2011), with organismal-group-specific pre-trained reference databases. For bacteria, fungi and metazoa, the non-redundant SILVA database (Glöckner et al., 2017) (*silva-138-ssu-nr99-seqs-derep-uniq*, version 138, 99% identity criterion) was pre-trained to generate amplicon-region specific classifiers. For the taxonomic assignment of protists, the pr2 reference database (Guillou et al., 2012) was used to build the pre-trained amplicon-region specific classifier. QIIME2 output files were imported in R as ‘phyloseq objects’ with the function *import_qiime* of the phyloseq package (v1.34.0) (McMurdie and Holmes, 2013). The phyloseq package was also used to process the phyloseq objects prior to the statistical analysis. Unassigned ASVs and ASVs assigned to chloroplasts and mitochondria as well as to non-target organisms were filtered out using the function *subset_taxa*. Samples with an unacceptably low number of reads (<1000) were filtered out using the *prune_samples* function. In the R package metagMisc (v0.0.4) (Mikryukov, 2017) the function *phyloseq_filter_prevalence* was used to filter out singletons and ASVs with less than 10 reads in the whole dataset.

### 2.5 Microbiome diversity and composition

ASV diversity and richness of each cover crop rhizosphere and fallow were determined for each organismal group using rarefied ASV tables. Rarefying was performed in the phyloseq R package using the function *rarefy_even_depth* with the options ‘with replacement’ and ‘to the minimum library size’. The function *alpha* of the microbiome R package (v.1.12.0) (Lahti, 2012-2019) was used to calculate Observed, Chao1 and Shannon diversity and richness metrics. In the case of normally distributed diversity scores, one way ANOVA and Tukey HSD post-hoc tests were used to pinpoint significant differences among cover crop species. Kruskal Wallis and Wilcoxon post-hoc tests with Bonferroni correction for multiple testing were employed for non-normally distributed alpha diversity scores.

Differences in microbial community structure across cover crops were calculated by constructing dissimilarity matrices with the Bray-Curtis Distance metric on the non-rarefied normalised ASV tables. Normalisation was carried out on the ASV tables using the Cumulative Sum Scaling (CSS) method (*cumNorm* function from R package metagenomeSeq v. 1.32.0) (Paulson et al., 2013). The results of the distance metrics were visualized in Principal Coordinate Analysis (PCoA) graphs for each organismal group, built with the function *plot_ordination* implemented in the phyloseq R package. The factors explaining the dissimilarities among the microbiomes of the different treatments were tested using PERMANOVA (*adonis2* function from the vegan package, Oksanen et al. (2013)). The factors included in the PERMANOVA were: subplot (to account for the block effect), nucleic acid (to account for the difference between DNA and cDNA), and treatment (to account for the effect of the cover crop treatments). Pairwise PERMANOVAs were carried out to calculate the comparisons between cover crop treatments and fallow and among cover crop treatments, with Benjamini-Hochberg correction for multiple testing. The R^2^ values associated with the pairwise PERMANOVA test were used as a proxy of the dissimilarity between treatments with high values indicating a higher dissimilarity and low values meaning a milder dissimilarity.

The taxonomic composition of the cover crops’ microbiome was visualised with stacked bar plots, generated using the function *plot_composition* of the microbiome R package. The plots show the relative abundance per cover crop treatment of the most abundant microbial families (bacteria and fungi) or orders (metazoa and protists), while taxa represented by less than 2% of all reads were grouped in the “Other” category.

### 2.6 Differential abundance analyses of microbial features in the rhizosphere using ANCOM-BC

A Differential Abundance analysis (DA) of microbial taxa was performed with Analysis of Compositions Of Microbiomes with Bias Correction (ANCOM-BC) in R (ANCOMBC R package) (Lin and Peddada, 2020). The test aimed to pinpoint the differential effect of each cover crop rhizosphere on the fallow soil. Cover crops species and block were the covariates of interest, while the fallow soil (control) was used as the reference level. The beta coefficients resulting from the linear regression indicated depletion (negative values) or enrichment (positive values) of the differentially abundant taxa in the rhizosphere of each cover crop compared to the fallow. Beta coefficients of the differentially abundant taxa were plotted in heatmaps, and associated dendrograms were generated based on Euclidean distances. ANCOM-BC was performed at the family level for bacteria and fungi, and at the order level for protists and metazoa.

## 3. RESULTS

### 3.1 Sequencing results

The sequencing of 74 DNA and 74 cDNA rhizosphere samples from ten cover crop species and the bulk soil (fallow) treatment resulted in a total of 24,823,255 reads from four organismal groups, i.e., bacteria, fungi, protists and metazoa. After pre-processing and filtering, 6,082,172 (2,886,425 DNA and 3,195,747 cDNA) reads were retained and used for further analyses, of which 3,252,176 belonged to bacteria, 868,312 to fungi, 1,475,652 to protists and 486,032 to metazoa, with median of 30,431 reads per DNA and 26,754 per cDNA sample. Samples featuring less than 1,000 reads were filtered out from each organismal group’s dataset (see also Table 3). Due to a high number of samples with a low number of reads in the metazoan cDNA dataset, the analyses of the metazoan community were based on DNA data only. 6,631 ASVs were assigned to bacterial taxa, 258 to fungal taxa, 1,812 to protistan taxa and 82 to metazoan taxa. For bacteria and fungi, the taxonomic resolution allowed for investigation of the communities up to the family level, whereas protists were mainly studied at the order level, as a minority of ASVs were assigned to lower taxonomic levels. The metazoan community was only studied at the order level.

### 3.2 Factors affecting the rhizosphere microbiome composition

The microbiome composition of the resident (DNA) and active (cDNA) microbial communities significantly differed. PERMANOVA showed that the factor ‘nucleic acid type’ explained 12% of the overall in both the bacterial and the fungal communities (p < 0.001), and 13% in the protist’s community (p < 0.001) (visualised in Suppl. Figure S1, Suppl. Table 5). Hence, active and resident microbial communities were analysed separately. The effect of blocks was significant (PERMANOVA < 0.001) but limited as it explained only 7.2, 6.4 and 8.4% of the variation across the bacteria, fungal and protistan communities. For the metazoan community, however, the block effect was more prominent, as it accounted for 17.9% of the overall microbiome variation (Suppl. Table 5).

Cover crop species had a major effect on the assembly of both the resident and active fractions of the rhizosphere microbiome for all four organismal groups, explaining 33 to 51% (PERMANOVA, p < 0.001) of the overall variation (Table 3, Fig. 1). The smallest effect was observed for the resident metazoan community, whereas the strongest impact was detected for the active fraction of the fungal community (Table 3). Both oilseed radish cultivars showed the strongest effect on microbial communities across all cover crop species, as shown by clearly separate data clusters in PCoA ordinations (Fig. 1). Contrastingly, the Poaceae crops black oat, hybrid ryegrass, and tall fescue induced relatively mild shifts as compared to the fallow control, as indicated by the relative proximity of samples in the PCoAs (Fig. 1). These patterns were supported by the PERMANOVA analyses, in which the highest dissimilarity values (based on R^2^) were associated with the rhizosphere of the two oilseed radish cultivars, and the lowest with Poaceae crops and marigold (Suppl. Table 6). These patterns were consistent across bacterial, fungal and protistan groups in both the active and resident communities (Suppl. Table 6). Deviating from this pattern, the fungal rhizosphere community of marigold differed greatly from the fallow fungal community. The resident metazoa community of borage was the most dissimilar from the fallow community, followed by oilseed radish, lentil and vetch.

**Figure 1.**
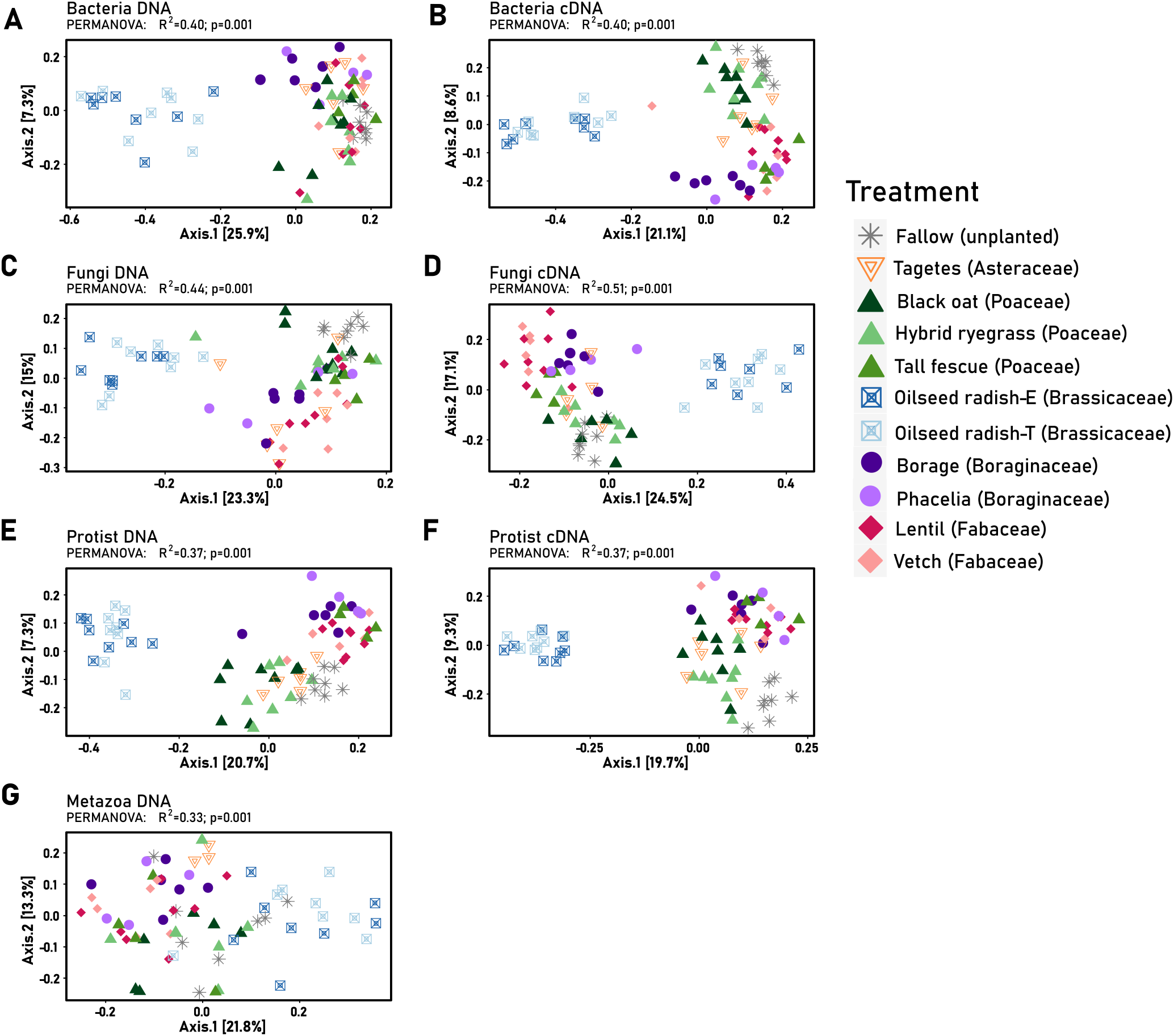
Principal Coordinate Analysis (PCoA) of CSS normalised ASV data. Dissimilarity matrix built on Bray-Curtis metric and plotted separating ASV based on cover crop treatment. Samples clearly appear separated along the axes as cover crop treatment accounted for between 19 and 26% of variation along the principal PCoA axis and explained 33 and 51% of the dissimilarity among sample groups (PERMANOVA, p ≤ 0.001).

### 3.3 Impact of cover crops on the microbial richness and diversity in the rhizosphere

To assess the effects of individual cover crops on the ASV richness and diversity, three diversity indices were used (Observed, Shannon and Chao1). All three indices identified vetch, tall fescue, lentil, phacelia and borage as cover crops associated with resident and active bacterial diversities that exceeded the diversity in the fallow controls. The two oilseed radish cultivars showed alpha diversities lower than the fallow controls (Suppl. Table 7-A, B).

As compared to the bacterial communities, the impact of cover crops on the richness and diversity of fungal assemblages in their rhizospheres was relatively mild. The four representatives of the families Fabaceae and Boraginaceae, vetch, lentil, borage and phacelia, as well as tall fescue (Poaceae) were associated with more diverse resident and active fungal communities compared to fallow by the indices Observed and Chao1. However, both for the resident and the active fractions, the Shannon index did not reveal significant differences between the fallow control and the aforementioned cover crops (Suppl. Table 7-C, D).

Also for the protists, vetch, lentil, borage and phacelia, as well as tall fescue were often associated with microbiomes showing a higher diversity than the fallow control. However, the Shannon index pointed to the absence of significant differences in the resident protistan communities between most cover crop species (except for black oat and the two oilseed radish cultivars) and the fallow control. Regarding the active protist fractions, the Shannon index did not reveal a significant difference between the fallow control and any of the cover crops (Suppl. Table 7-E, F). Concerning the resident metazoan community, borage and vetch were associated with the most diverse resident community (Suppl. Table 7-G).

Analyses of the alpha diversities in the rhizospheres of ten cover crop species as compared to the fallow controls using three indices revealed significant negative as well as positive changes in the bacterial resident communities as well as the active fractions thereof. It is noted that changes in the alpha diversity of the resident communities relate to the detectability of individual taxa. Cover crops may stimulate rare microbial taxa. As a result, they became detectable, and give rise to an alpha diversity increase. Regarding the other organismal groups, fungi, protists and metazoa, the effect of the cover crops on the alpha biodiversity was minor.

### 3.4 Impact of cover crops on the relative microbial composition in the rhizosphere

In Fig. 2 the relative microbial composition of the rhizosphere is shown for each of the ten cover crop species, as well the composition of the fallow control. The impacts on each of the four organismal groups are presented, and panels showing the composition of the resident and the active community are displayed next to each other. A selected number of results will be detailed below.

**Figure 2.**
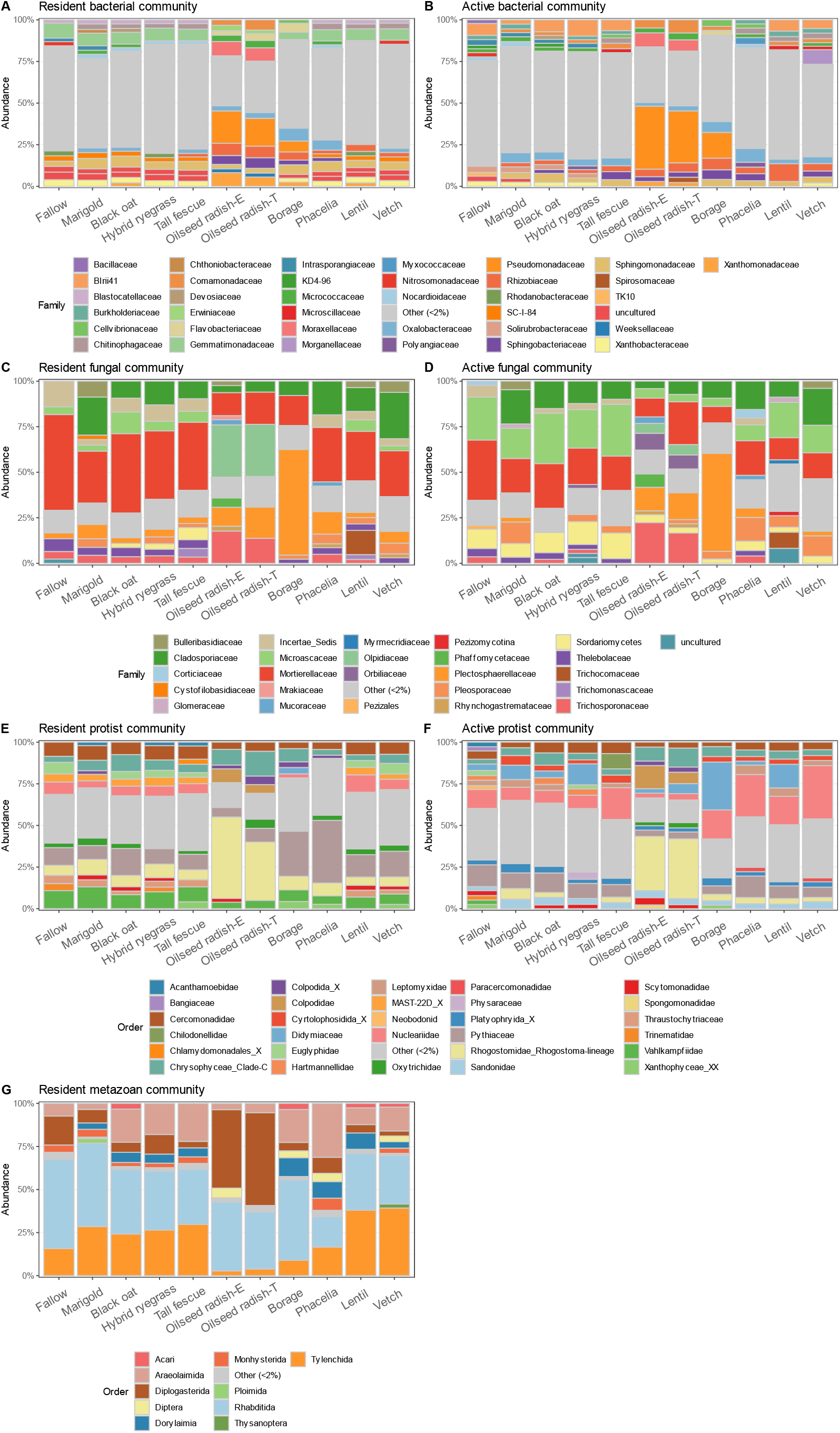
Microbial composition at family level of the rhizosphere of the cover crops. A) bacterial resident community, B) bacterial active community; C) Fungal resident community; D) fungal active community; E) protists resident community; F) protists active community; G) metazoa resident community(order level). All taxa accounting for <2% or relative abundance were grouped as “Others (<2%)”.

#### Bacteria

Numerous rare bacterial taxa (‘other (<2%)’ in Fig. 2) were present in the rhizosphere of all cover crops tested. In the case of hybrid ryegrass, tall fescue and lentil, this category comprised >50% of both the resident and the active bacterial community. Among the bacterial families with abundances >2%, striking differences were observed between the individual cover crop species. Pseudomonadaceae were highly enriched in the rhizosphere of the two oilseed radish cultivars (on average 18.1% and 34.6% of the resident and active community respectively) and borage (6.6% and 15.5% of the resident and active community respectively) (Fig 2A, B). Moreover, Moraxellaceae and Rhizobiaceae were abundantly present in the rhizospheres of the two oilseed radish cultivars. On average Moraxellaceae accounted for 8.4% of the resident and 7.6% of the active community, whereas the Rhizobiaceae were amply represented in the resident community (7.4%). Gemmatimonadaceae (≈7%) and Xanthobacteraceae (2.4-3.6%) were well represented in the resident bacterial communities of all cover crops, except for oilseed radish (both cultivars). Notably, the Gemmatimonadaceae comprised <2% of the active fractions for all cover crops (Fig. 2-B). Regarding the active fraction of thebacterial communities, Blrii41 (order Polyangiales) was most abundant in the hybrid ryegrass rhizosphere (9.8%). This bacterial family was activated by cover crops belonging to Poaceae, Asteraceae and Fabaceae, and not by representatives of the Brassicaceae and Boraginaceae. It is noted that Oxalobacteraceae were activated by all cover crops, while the active members of this bacterial family made up <2% in the fallow control (See Suppl. Table 8 for details on the composition of the bacterial communities).

#### Fungi

As compared to bacteria, the fraction ‘other (<2%)’ is relatively small in the fungal communities (Fig. 2-C, D). The fungal family Mortierellaceae was abundantly present in the rhizospheres of all cover crops tested, as well as in the fallow control. The relative abundances of this fungal family ranged from 9.2% (active community of borage) to 53% (resident community of fallow control). Notably, the representation of this fungal family in the resident community tended to exceed its relative presence in the active community. An opposite pattern was observed for the Microascaceae; although present within the resident community of most cover crops, they constituted a larger part of the active fraction of the fungal community. At the level of individual cover crops, the rhizosphere of borage showed the highest abundance of Plectosphaerellaceae (58.1% and 53.8% in the resident and active community respectively) (p < 0.05). The two oilseed radish cultivars stood out as their rhizospheres were highly enriched in Olpidiaceae (on average 28.7% of the resident community). It is noted that the presence of this fungal family was much lower in the active fraction (on average 5.5%). An opposite trend was observed for the Orbiliaceae, active representatives of this fungal family were amply present in the rhizosphere of the two oilseed radish cultivars (on average 8.5%), while the relative abundance in the resident community was <2% (See Suppl. Table 9 for details on the composition of the fungal communities).

#### Protists and Metazoa

The resident and active protistan communities showed a high relative abundance of rare taxa (Fig. 2-E, F, category ‘other (<2%)’). Among the protist taxa present at higher relative abundances (>2%), the Cercomonadidae (order Cercomonadida) and Pythiaceae (order Peronosporales) were shown to be present and active in the rhizospheres of all cover crops tested. With relative abundances of 27.1% and 37.7% in the resident communities of borage and phacelia, the oomycete family Pythiaceae was dominantly present in the rhizosphere of Boraginaceae (Fig. 2-E, F). Pythiaceae were also detected in the active fractions, though at lower relative abundances. In the case of the two oilseed radish cultivars, the Rhogostomidae (order Cryomonadida) were remarkably well represented in both the resident (on average 40.6%) and the active (on average 35.5%) fraction of the protistan community. The Vahlkampfiidae were abundant in the resident community of all cover crops rhizosphere, but they made up less than 2% of the active protist fraction. An opposite trend was observed for the Nucleariidae and Sandonidae, which were abundant in the active community all cover crops rhizosphere, but underrepresented in the resident community.

Analysis of the resident metazoan community in the rhizosphere of cover crops revealed the dominance of nematodes (seven of the most abundant orders shown in Fig. 2-G). Rhabditida (17.7-46%), opportunistic bacterivores, were abundantly present in the rhizosphere of all cover crops. Diplogasterida (bacterivores) were present in remarkably high abundances in the rhizosphere of the two oilseed radish cultivars (on average 50%) as compared to the other cover crops (2-9%). Representatives of the orders Tylenchida and Areolaimida were commonly present in all cover crops but less so in the two oilseed radish cultivars (∼3% *versus* ∼20-30%) (See Suppl. Table 10 for details on the composition of protist and metazoan communities).

### 3.5 Differential abundance analysis of the rhizosphere microbiomes with ANCOM-BC

In the next step, an analysis of compositions of microbiomes with bias correction (ANCOM-BC, Lin et al. 2020) was used to determine differential abundances for each of the four organismal groups per cover crop as compared to the fallow soil controls. Beta coefficients, a quantitative measure for differential abundance, ranged from -4.4 (Microascaceae under oilseed radish) to 7.2 (Pseudomonadaceae under oilseed radish) indicating that, overall, the stimulation of taxa in the rhizosphere by the cover crops was stronger than the repression. As this analysis concentrates on changes in abundances rather than abundances *per se*, the heatmaps generated with the beta-coefficient values include taxa present in low abundance that were lumped under the category ‘other (<2%)’ in Fig. 2. On the other hand, taxa that were shown to be present in the rhizosphere of all cover crops in relative abundances comparable to fallow, such as the bacterial family Sphingomonadaceae, are not included in Fig. 3.

**Figure 3.**
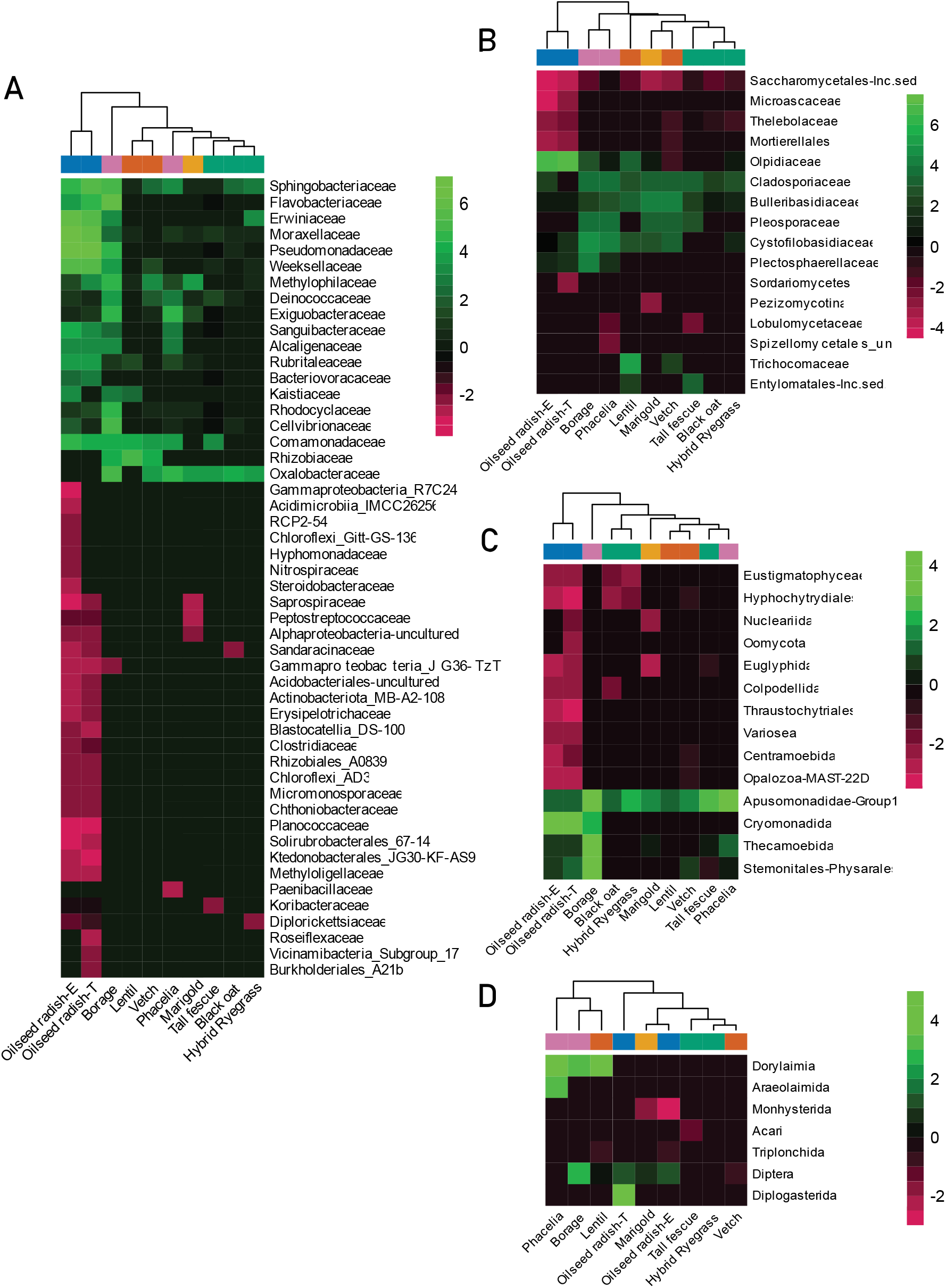
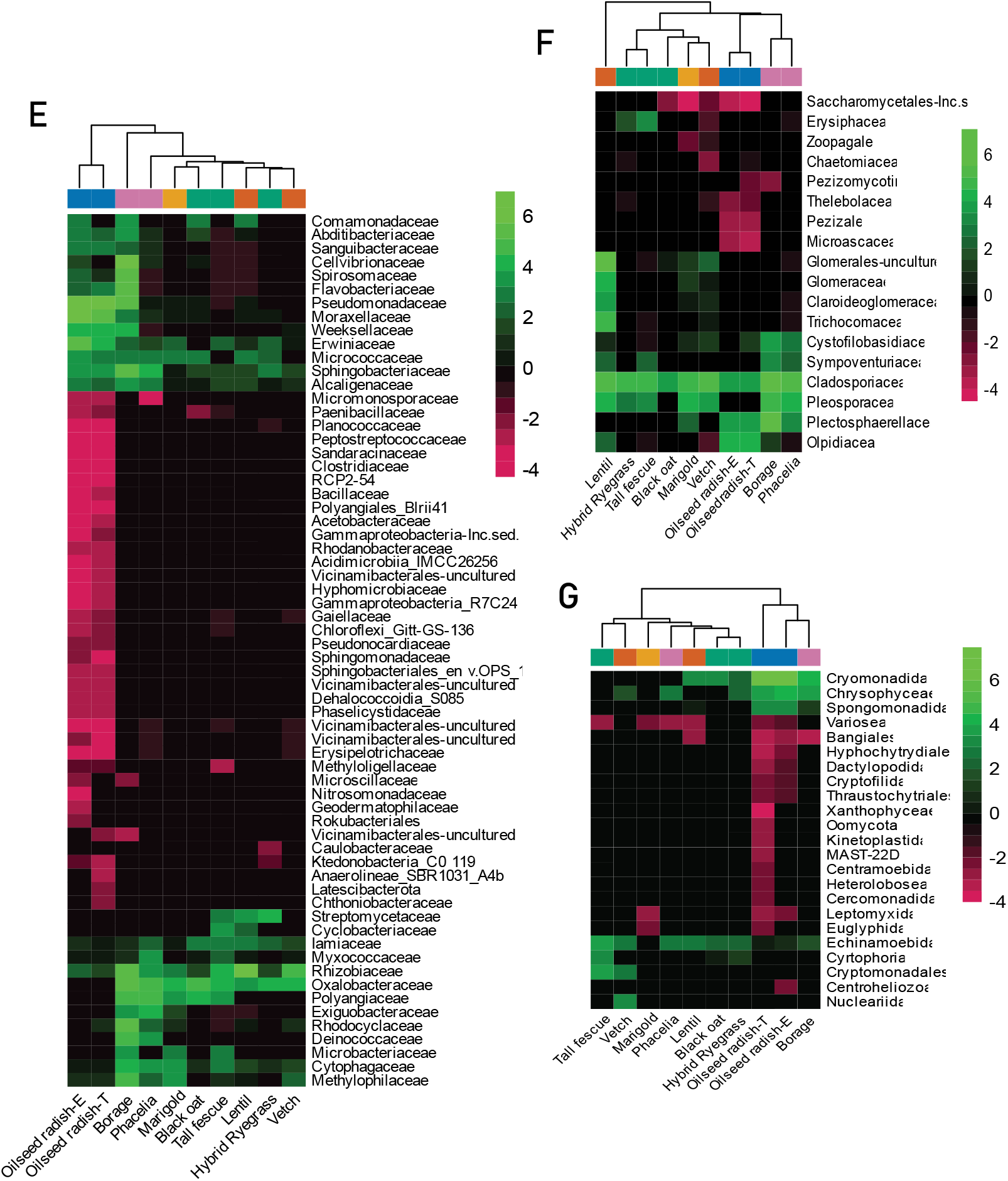
Heatmap of the beta coefficient assigned to the differentially abundant taxa pinpointed with ANCOM-BC for A) resident bacterial families, B) resident fungal families, C) resident protist orders, D) resident metazoan orders, E) active bacterial families,, F) active fungal families,, G) active protists orders,. Beta coefficients greater than zero (shades of green) indicate that the average abundance of the taxon in the cover crop treatment is higher than in the reference level (fallow soil). Beta coefficients lower than zero (shades of red) indicate that the average abundance of the taxon in the cover crop treatment is lower than in the reference level (fallow soil). The dendrograms per column and row were calculated based on Euclidean distance. The colours above the heatmap represent the plant family of each cover crop cultivar: orange = Asteraceae, green = Poaceae, blue = Brassicaceae, purple = Boraginaceae, and red = Fabaceae. For improved figure readability, for bacteria, fungi and protists only taxa with beta coefficients higher than 3 and lower than -2 are represented. The complete heatmaps are available as Supplementary Figure S2.

#### Bacteria

ANCOM-BC identified 175 and 177 differentially abundant families among the resident and active bacterial communities, respectively (a selection of the most enriched and most depleted families is presented in the heatmap in Fig. 3 - A, E, the complete heatmap representing all the differentially abundant families is provided as Suppl. Figure 2 - A, D). Of these, 69% (120) was shared between the two, indicating they showed differential abundances in both the resident and active community fractions (see Suppl. Table 11). The two oilseed radish cultivars resulted in bacterial profiles that were most deviant from the fallow controls. Despite a few cultivar-specific changes both in the resident and active communities, the two oilseed radishes clustered together in dendrograms based on Euclidean distances (Fig. 3-A, E). In the rhizosphere of the oilseed radishes, the number of repressed bacterial taxa exceeded the number of promoted taxa for both the resident and the active bacterial communities. Planococcaceae (both in resident and active communities) and Bacillaceae (in the active community) were among the most repressed families, while Pseudomonadaceae, Moraxellaceae and Erwiniaceae resided among the most enriched families (both in resident and active communities). A relatively high number of repressed families was also observed in the resident community of marigold and the active bacterial fraction of tall fescue (Fig. 3-A, E). Borage showed a bacterial profile exceptionally distinct from the fallow soil, with a high number of differentially enriched families (Fig. 3-A, E). Among these, the resident and active fractions of the Flavobacteriaceae, Cellvibrionaceae and Sphingobacteriaceae were strongly promoted as compared to fallow controls (in all cases beta coefficients > 5).

#### Fungi

ANCOM-BC pinpointed 44 and 46 differentially abundant fungal families in respectively the resident and active communities of the cover crop rhizospheres (see Suppl. Table 12 for ANCOM-BC results on the fungal community, and Suppl. Figure 2-B, E, F for the full heatmap). The resident and active fungal communities shared a high number of differentially abundant taxa (35 families, about 78%). Nevertheless, remarkable contrasts were observed between the resident and the active fractions of the fungal communities (Fig. 3-B, F). Regarding the resident community, the two oilseed radish cultivars showed a high number of differentially abundant fungal families. Most striking are the strong repression of the representatives of the bacterial family Microascaceae and the order Saccharomycetales, and the stimulation of members of the protistan family Olpidiaceae (all both in the resident and active community). Apart from a mild repression of Saccharomycetales (beta coefficient of -1.55), the impact of lentil on members of the fungal community was invariably positive (Fig. 3-B, F). The stimulating effect was most notable for the fungal family Trichocomaceae (in the resident and active communities) and the fungal order Glomerales (in the active community). In Fig. 3-F, this order is represented by ‘Glomerales-uncultured’, ‘Glomeraceae’ and ‘Claroideoglomeraceae’. For both representatives of the Boraginaceae, borage and phacelia, stimulation of the Cystofilobasidiaceae and the Pleosporaceae were observed both in the resident and the active fraction of the fungal community (Fig. 3-B, F). It is noted that due to its low relative abundance, the fungal family Cystofilobasidiaceae resides under ‘other (<2%)’ for borage in Fig. 2-C, D. The fungal family Cladosporiaceae is exceptional as its members were stimulated by virtually all cover crops in both the resident and active communities (only exception oilseed radish, cultivar -T) (Fig. 3-B, F).

#### Protists and metazoa

As main consumers of primary decomposers, protists and metazoans were indirectly affected by cover crops. ANCOM-BC analyses revealed that 46 and 48 protist orders were differentially abundant in the resident and active rhizosphere communities, respectively. Thirty-six orders were affected at both DNA and cDNA levels. The two oilseed radish cultivars had the broadest impact on the resident and active protistan communities, followed by borage (Suppl. Table 13 and Suppl. Figure 2-C, F for the complete heatmap of the protistan community). Cryomonadida was relatively most enriched in the oilseed radish and borage rhizosphere compared to fallow. Also enriched in borage were the resident members of the Stemonitales-Physarales (plasmodial slime moulds). It is noted that these protist orders were present in low abundances (<2% in Fig 2). ANCOM-BC of the metazoan community identified 7 differentially abundant metazoan orders out of 79 (Suppl. Table 11). The highest number of differentially abundant metazoan orders was found for marigold (Fig. 3-D). The nematode order Diplogasterida was exclusively enriched in the rhizosphere of the oilseed radish cultivar-T, and Monhysterida was depleted in oilseed radish cultivar -E and marigold, while Araeolaimida were enriched in the rhizosphere of phacelia (Figure 3-G). It is noted that the sample size (2 g of rhizosphere soil) is low for metazoa, and the shifts reported here require confirmation by the analysis of more and larger subsamples.

## 4. DISCUSSION

### 4.1 Cover crop species exert different selection strengths on the rhizosphere microbiome

All ten cover crops characterized in this study exerted significant effects on the soil rhizobiome, and the kind of effects was shown to be plant species and - to some extent - plant family dependent. The characterization of the resident and active fractions of the rhizosphere communities of the cover crops revealed three distinct levels of selection strengths by the cover crops: strong, intermediate, or mild.

The highest selection strength in the rhizosphere was brought about by the two cultivars of oilseed radish. Both oilseed radish cultivars sharply suppressed a wide range of microbial taxa, in particular bacterial and protistan microorganisms, and, at the same time, strongly promoted a smaller subset. It is noted that members of the Brassicaceae plant family, including oilseed radish, produce a category of secondary metabolites called glucosinolates. The release of these metabolites and their biocidal hydrolysis products in the rhizosphere by living roots was demonstrated for canola (Choesin and Boerner, 1991) and mustard roots (Schreiner and Koide, 1993) and directly impacted the rhizobiome (Bressan et al., 2009). We suggest that the release of glucosinolates and their breakdown products might have contributed to the observed high selection strength of both oilseed radish (*Raphanus sativus* var. *oleiformis*) cultivars. With a strong promotion of both the resident and the active fractions for multiple bacterial and fungal families, borage resembled the oilseed radish cultivars. As its suppressing selection strength is milder than observed for the oilseed radishes, borage should be seen as a less pronounced representative of this category.

A diverse set of cover crop species including phacelia (another species representing Boraginaceae), vetch and lentil (both legumes) and tall fescue (Poaceae) imposed an intermediate selection strength on the rhizosphere microbiome following different patterns. As compared to borage, phacelia showed overall a milder impact on the microbiome. Especially the active fraction of the fungal community was less strongly affected by phacelia than borage. Lentil showed a high number of differentially enriched fungal taxa in its rhizosphere, while no significant depletion was shown for any fungal taxon compared to fallow soil. Similarly, tall fescue featured a prevalence of differentially enriched microbial taxa in its rhizosphere and a minority of differentially depleted ones.

Cover crops with a mild effect on the rhizosphere microbiome included two Poaceae species - black oat, hybrid ryegrass - and marigold (Asteraceae). It is noted that for marigold, the selection strength for one of the organismal groups - the fungal community - was comparable to the investigated cover crops with an intermediate selection strength. Two out of the three Poaceae cover crop species were shown to have mild selection strength on their surrounding soil microbiome. Black oat belongs to the subtribe Aveninae, while hybrid ryegrass and tall fescue are phylogenetically more related (both reside in the subtribe Loliinae). Hence, there is no phylogenetic rationale that could explain why tall fescue had a stronger impact on its rhizobiome than the two other Poaceae cover crop species. However, tall fescue tends to have thicker and deeper root systems as compared to ryegrasses (Cheng et al., 2016). Because root traits impact the release of root exudates (Saleem et al., 2018), this may explain why tall fescue exerted a selection strength that differed from cover crops belonging to the same plant family.

Distinctions in selection strengths on the soil microbiome between plant species have been reported before. In a comparison between three model plants and three crops, *Arabidopsis* was shown to exert the weakest influence on its rhizobiome (Tkacz et al., 2015). In a comparison between *Festuca arundinacea, Cichorium intybus* (chicory) and legume *Medicago sativa, C. intybus* was shown to have the strongest impact on the bacterial community (Uksa et al., 2014), and recently, Cloutier et al. (2021) demonstrated that soybean has a stronger selective effect on bacterial-archaeal assemblages than corn. To the best of our knowledge, the categorisation of a range of cover crops based on their microbial selection strengths is new.

### 4.2 Compositional changes underlying differential selection strengths

Differences in selection strengths are reflected in specific shifts within the bacterial, fungal, protist or metazoan community. Dozens of individual taxa were shown to be affected by cover crops and here we will highlight five of them. Three examples involve soil microorganisms that were shown to be promoted by at least one of the cover crops investigated here, while two examples are given of cover crop-repressed microorganisms.

The increased abundance of the Pseudomonadaceae bacterial family in the rhizosphere of some cover crops was remarkable. This bacterial family constituted up to 37.9% and 15.5% of the oilseed radish and borage bacterial community in the rhizosphere and yielded beta coefficients up to 6.9. Strong plant-induced stimulation of the bacterial family Pseudomonadaceae has been reported before. Application of Brassicaceae seed meal in an apple orchard resulted in an increased abundance of *Pseudomonas* representatives (Wassermann et al., 2017; Wang and Mazzola, 2019). In a field experiment with four crops including canola (*Brassica napus*), this *Brassica* species was shown to strongly stimulate endophytic *Pseudomonas* representatives over multiple years and locations (Cordero et al., 2020), and - to a lesser extent - Pseudomonadaceae in the rhizosphere. The bacterial family Pseudomonadaceae harbours plant pathogens, beneficial species that can act as biological control agents as well as plant growth-promoting rhizobacteria (*e*.*g*., Lugtenberg and Kamilova (2009)). although the agronomical impact of the observed stimulation is currently hard to assess, the Pseudomonadaceae family is worthwhile to monitor as it has the potential to improve plant growth and health (Kim and Anderson, 2018).

Two arbuscular mycorrhizal fungal (AMF) families, Glomeraceae and Claroideoglomeraceae, showed an increased presence and/or activity in the rhizosphere of some of the cover crops tested. Previously, the presence of Claroideoglomeraceae was linked to mechanically disturbed soils, while Glomeraceae were more abundant in undisturbed habitats (Moora et al., 2014). The cover crops characterized here were grown in mechanically disturbed soil. Lentil, and to a lesser extent marigold and vetch, exclusively induced an enrichment of Claroideoglomeraceae. However, at the cDNA level, we observed an increased activity of both Glomeraceae and Claroideoglomeraceae for lentil, and less prominently for vetch and marigold. Our results do not contradict the results of Moora et al. (2014) as this study focused on Glomeromycota and was performed at the DNA level only. There was no AMF signal to be expected for the Brassicaceae (non-host for AMF (Cosme et al., 2018)), but we cannot explain why no increased AMF presence or activity compared to fallow soil was observed in the rhizosphere of the Poaceae included in this study.

The fungal family Cladosporiaceae showed an elevated presence in the rhizobiome in nine out of the ten cover crops tested, but most striking was the increase in its activity. This was most explicitly observed for borage, phacelia, lentil and vetch (beta coefficients > 5). The fungal family Cladosporiaceae harbours seven genera, the genera *Davidiella* and *Cladosporium* being by far the most widespread ones. *Davidiella* is most often found on aboveground tissues (Longley et al., 2020), whereas *Cladosporium* representatives are present both above- and belowground (Bensch et al., 2012). The genus *Cladosporium* harbours 189 described, mostly saprophytic, species (Sandoval-Denis et al., 2016a), and next to saprobes, this genus comprises above- and belowground endophytes and plant pathogens. Uncharacterized *Cladosporium* members were recently detected in the rhizobiome of maize (Zhao et al., 2021) and also – in a non-agricultural setting – in the rhizosphere of giant goldenrod (Harkes et al., 2021). We hypothesize that soil-borne, saprophytic and/or pathogenic *Cladosporium* species are responsible for the observed increased presence and activity in the rhizosphere of nearly all cover crops rhizosphere compared to fallow.

The bacterial family Planococcaceae was identified as the most strongly repressed bacterial family, and the repression was almost exclusively observed in the oilseed radish rhizospheres (next to a mild repression by hybrid ryegrass). The family Caryophanaceae/Planococcaceae is a polyphyletic bacterial family with >100 species classified within 13 genera (Gupta and Patel, 2020). Recently, *Planococcus* was observed as an endophyte in sugar beet (*Beta vulgaris*) roots and in higher relative abundances in its rhizosphere (Li et al., 2020). However, no ecological explanation could be given for this shift. The absence of any known characteristics exclusive to all bacterial members of the family *Planococcaceae* (Gupta and Patel, 2020) makes it impossible to assess the ecological impact of *Planococcaceae* in the rhizosphere.

With high beta-coefficients, Microascaceae belonged to the strongest repressed fungal taxa among all cover crops, and this repression was only observed for oilseed radish. Microascaceae currently accommodate a morphologically heterogeneous group of fungi, comprising saprophytic and plant pathogenic species (Sandoval-Denis et al., 2016b). Fungal members of the Microascaceae family inhabit niches in association with different kinds of bark beetles, *Petriella* and *Petriellopsis* are associated with soil, dung and compost (Lackner and de Hoog, 2011). We hypothesise that the toxicity of isothiocyanates associated with Brassicaceous plants may suppress this fungal family in the oilseed radish rhizosphere.

### 4.3 The inclusion of primary consumers in the characterisation of cover crops rhizosphere

The bacterial and the fungal communities in the cover crop rhizospheres are the resultant of the composition of the local microbial community in the soil, bottom-up control by the individual cover crops, and top-down regulation by primary consumers (Gao et al., 2019). The primary consumer community is here represented by protists and metazoa (mainly nematodes). It is noted that the primary consumer activity affects plant growth directly as grazing of the bacterial biomass by protists results in the release of plant-available N (Clarholm, 1985; Xiong et al., 2020). Essentially the same holds for the impact of selective grazing by bacterivorous nematodes (Schratzberger et al., 2019). Large shifts were observed both in the resident and the active fractions of the protist’s community. One of the most striking examples is the activation of the protist order Cryomonadida (Cercozoa phylum) in the rhizosphere of the two oilseed radish cultivars. It is premature to couple the increased activity of members of this predatory protistan order with the strong stimulation of Pseudomonadaceae in this cover crop species. However, a novel study by Guo et al. (2022) connected the enrichment of protistan predators with the stimulation of disease-suppressive bacteria belonging to *Bacillus spp*. in the rhizosphere. Further research is needed to determine whether this kind of association may also hold for members of the Pseudomonadaceae family and protistan predators.

Regarding the metazoa, it should be mentioned that the amount of soil analysed in this study, 2 g, might have been too small to get a proper representation of the metazoan community. A more complete representation of the nematode community would require an upscaling of the current DNA and RNA extraction procedure (Harkes et al., 2019).

## 5. CONCLUSIONS

Here we pinpointed the differential effects of ten cover crop species on both the resident and the active fractions of bacterial, fungal and protist communities in the rhizosphere. The ten cover crops under investigation had in common that the strength by which individual microbiome taxa are promoted exceeded the level at which other microbiota were suppressed. Our results prompted a classification of the cover crops under investigation into three main categories characterized by the exertion of mild, intermediate or strong selection strengths on the rhizosphere microbiome. Our data identified cover crops belonging to the families Poaceae and Fabaceae as being potentially suitable for a general stimulation of soil life. On the other hand, cover crops belonging to the families Boraginaceae, and, most explicitly, Brassicaceae could be used to steer the soil microbiome in distinct directions, *i*.*e*., by promoting a limited subset of microbial taxa. The current data set should be seen as a starting point for the application of specific cover crop species or mixtures thereof to steer the soil microbiome in a predictable direction to promote soil health and sustain healthy crop growth. Further studies should be aimed at determining to what extend the effects of cover crops on the soil microbiome persist over time and thus may affect growth and development of the main crop.

## 6. DATA AVAILABILITY

All sequences have been submitted to the NCBI database under BioProject ID PRJNA842568.

## 7. ACKNOWLEDGEMENTS

The authors would like to acknowledge the companies Barenbrug, DSV-zaden, Joordens Zaden and Vandinter Semo for their help in selecting and providing the cover crops seeds used in the experiment; Marc Kroonen, Harry Verstegen and the Vredepeel field team for setting up and maintaining the field experiment; Leendert Molendijk for the help in the experimental design. We also like to thank Joris van Steenbrugge and Lia Obinu for their help in the setting up of the bioinformatic pipelines; Paula Harkes for the guidance in the laboratory, Martijn Holterman, Francesco Garassino and Marco Albiero for helping with the soil sampling.

## SUPPLEMENTARY FIGURES

Supplementary figure S1 | Principal Coordinate Analysis (PCoA) of CSS normalised ASV data. Dissimilarity matrix built on Bray-Curtis metric and plotted separating ASV based on nucleic acid type (DNA vs cDNA). Samples appear separated along the second axis as nucleic acid accounts for between 13 and 17% of variation along the principal PCoA axis and explained 6 and 12% of the dissimilarity among sample groups (R^2^ PERMANOVA, p ≤ 0.001).

Supplementary figure S2 | Heatmap of the beta coefficient assigned to the differentially abundant taxa pinpointed with ANCOM-BC for A) resident bacterial families, B) resident fungal families,, C) resident protist orders, D) resident metazoan orders, E) active bacterial families, F) active fungal families, G) active protists orders. Beta coefficients greater than zero (in green) indicate that the average abundance of the taxon in the cover crop treatment is higher than in the reference level (fallow soil). Beta coefficients lower than zero (magenta) indicate that the average abundance of the taxon in the cover crop treatment is lower than in the reference level (fallow soil). The dendrograms per column and row were calculated based on Euclidean distance. The colours above the heatmap represent the plant family of each cover crop cultivar: orange = Asteraceae, green = Poaceae, blue = Brassicaceae, purple = Boraginaceae, and red = Fabaceae.

