## Supplementary figures and images for "Pinpointing the distinctive impacts of ten cover crop species on the resident and active fractions of the soil microbiome"

### Supplementary Figure 1

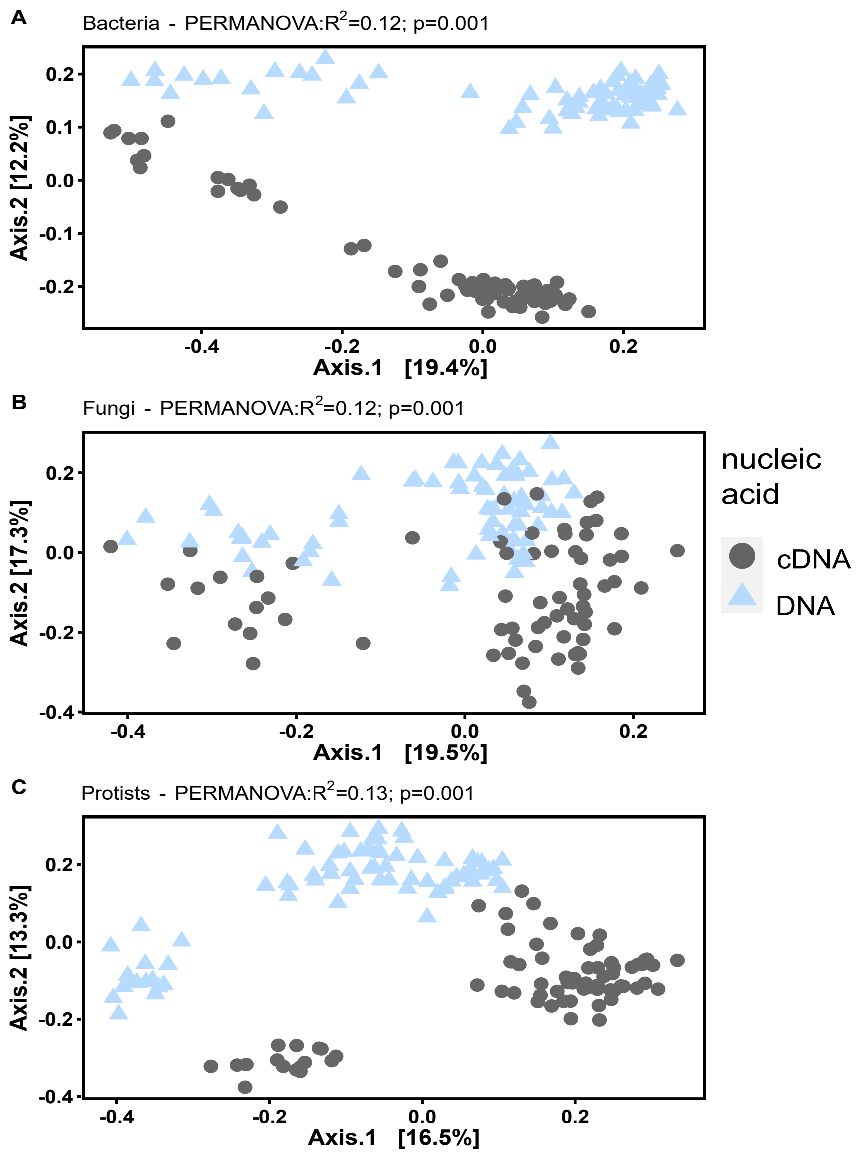

### Supplementary Figure 2

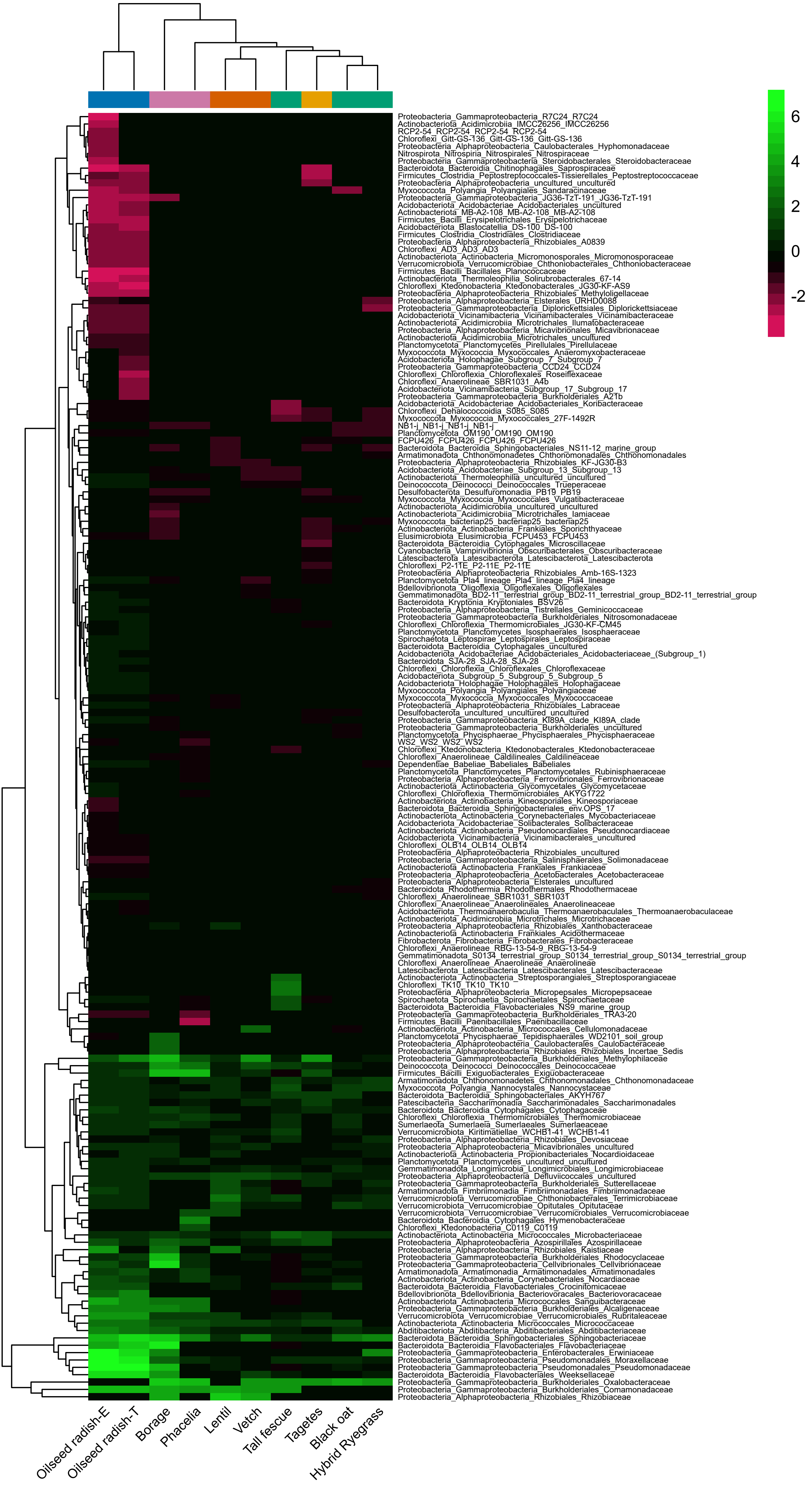

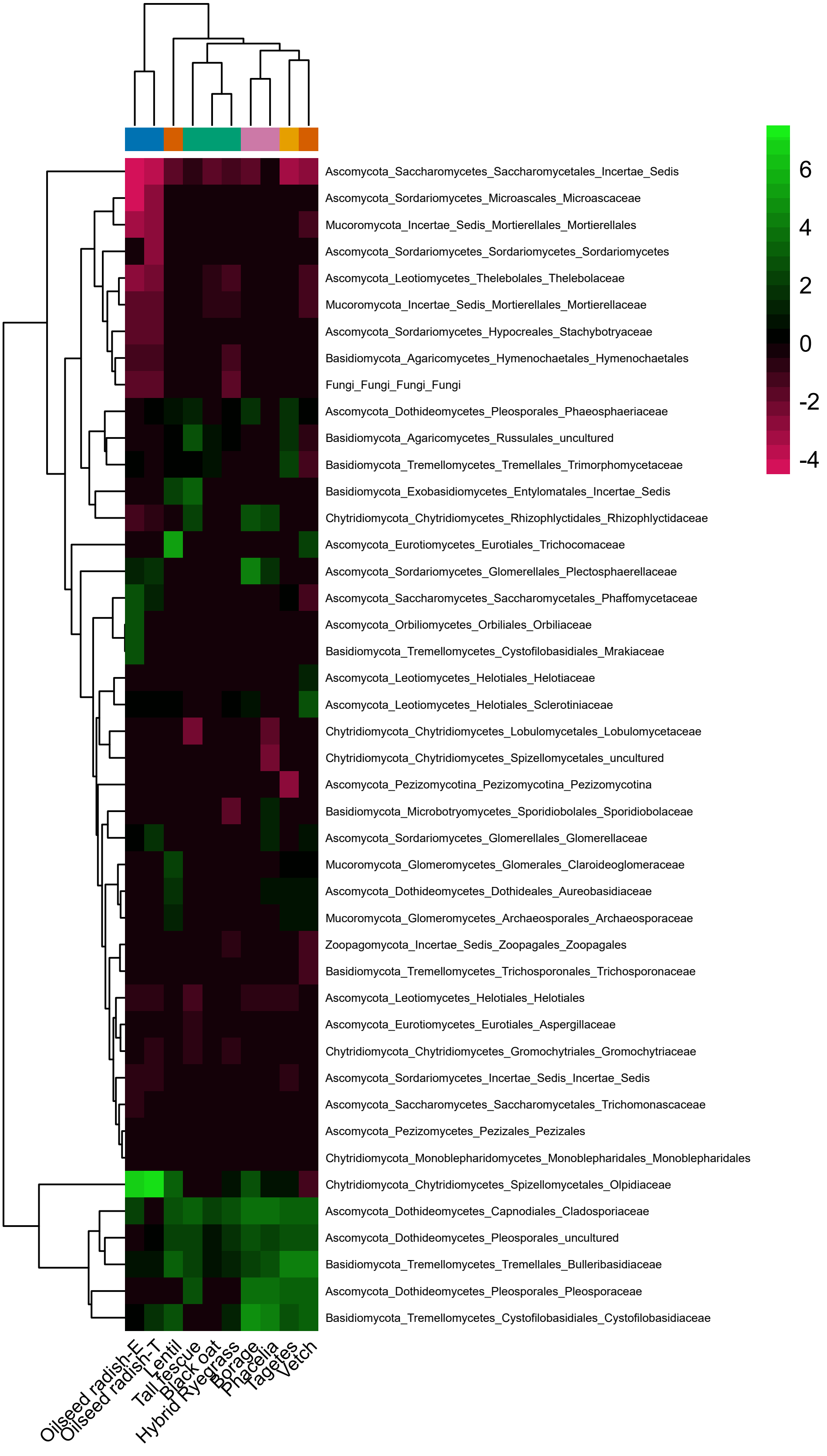

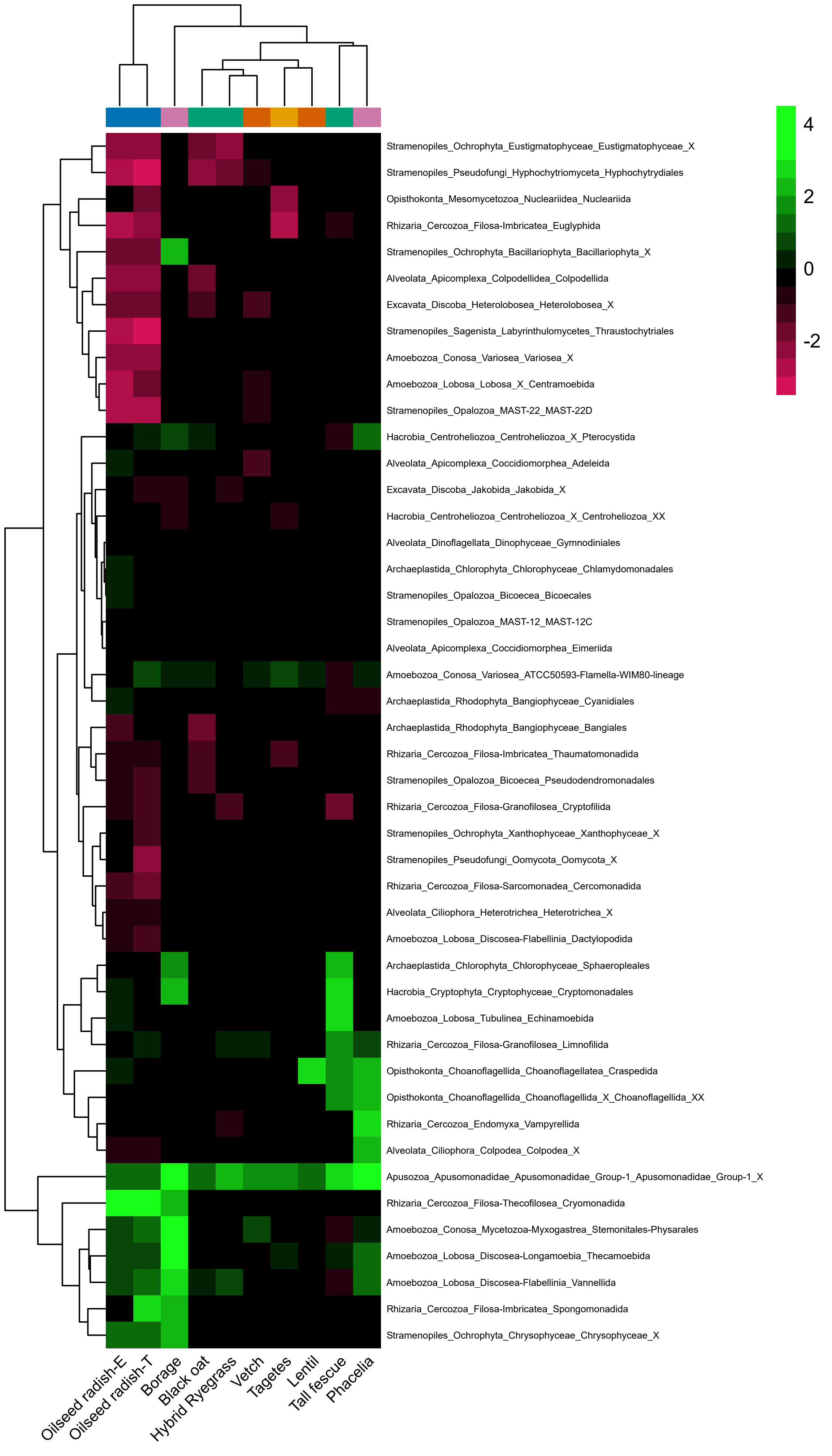

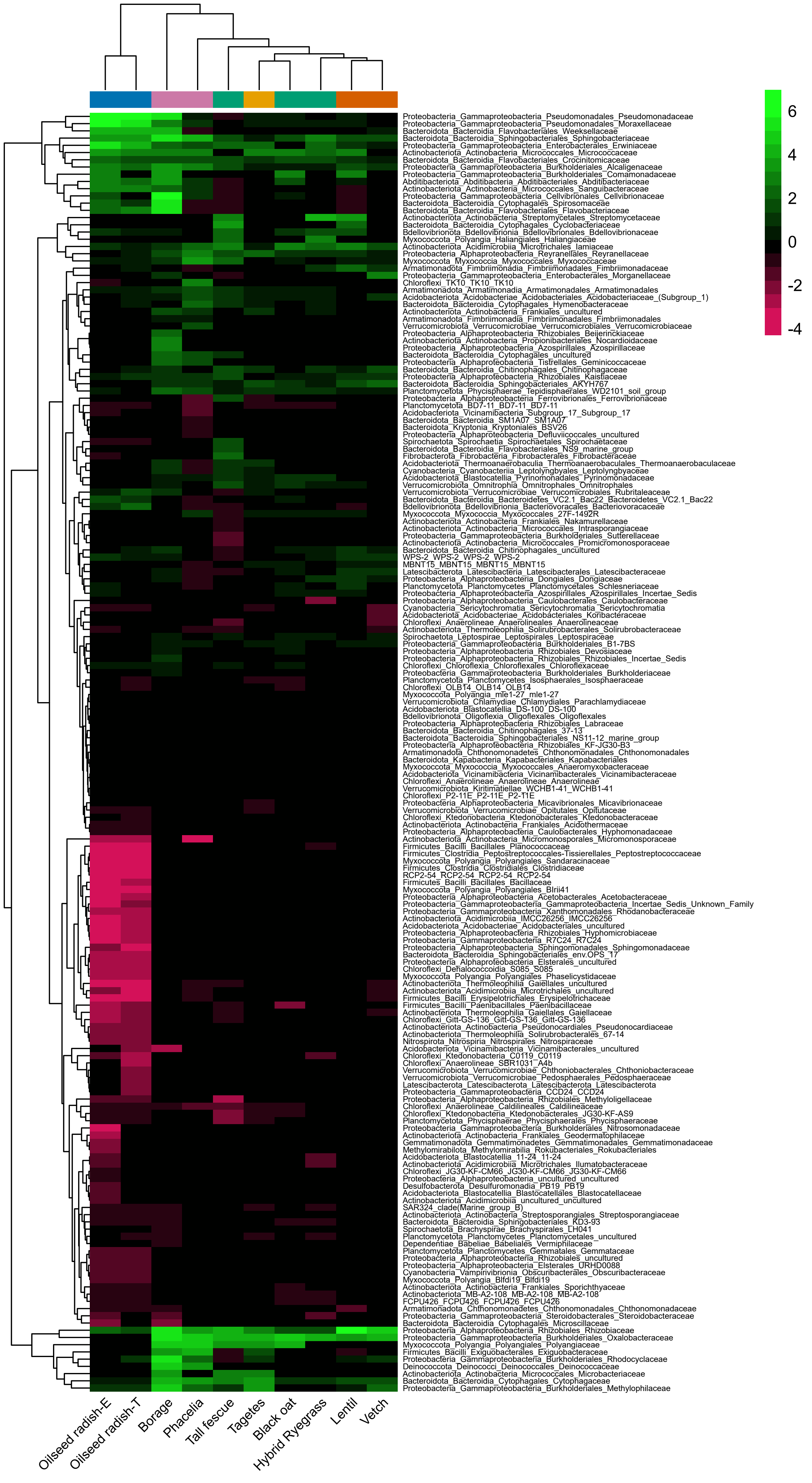

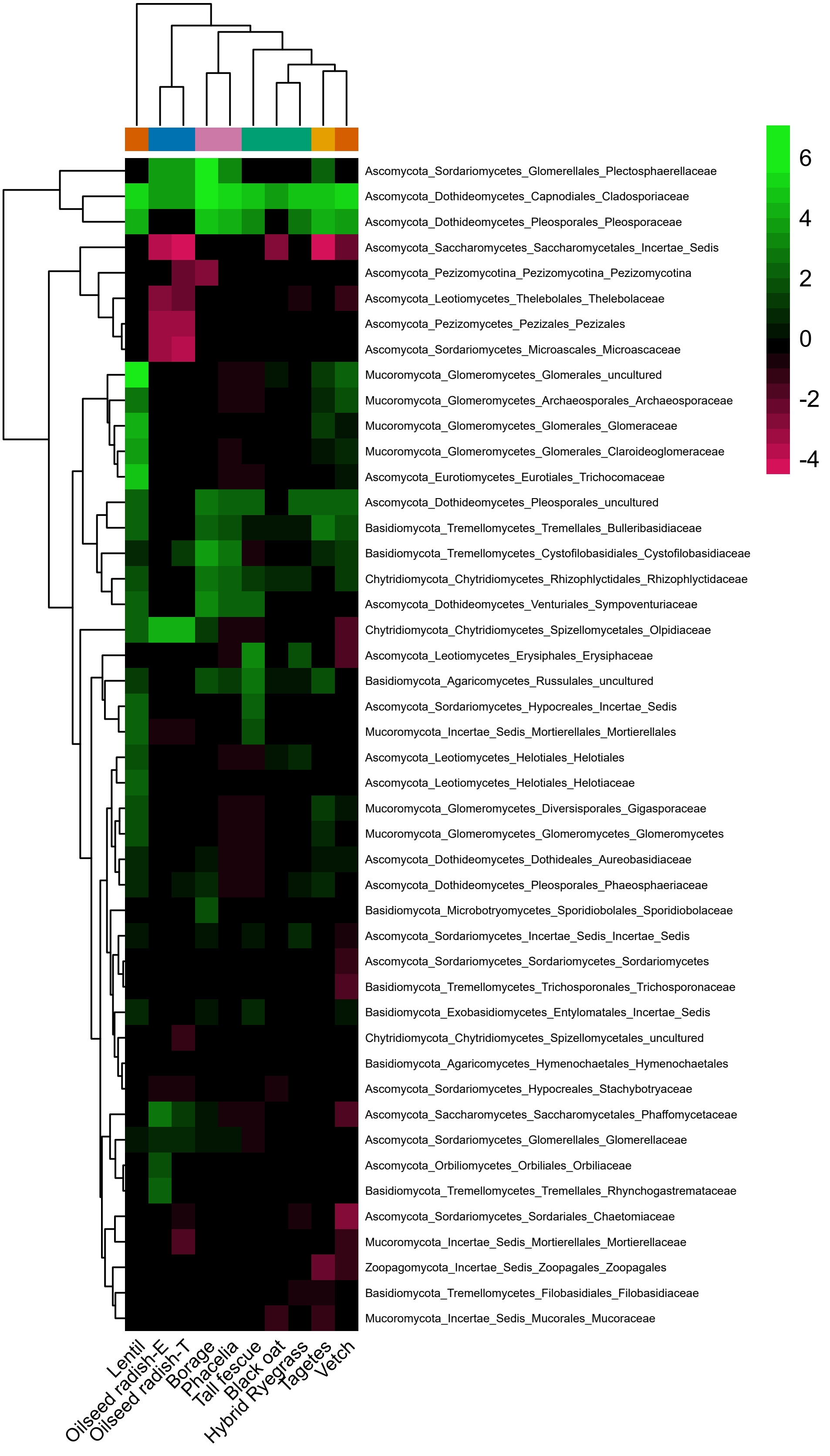

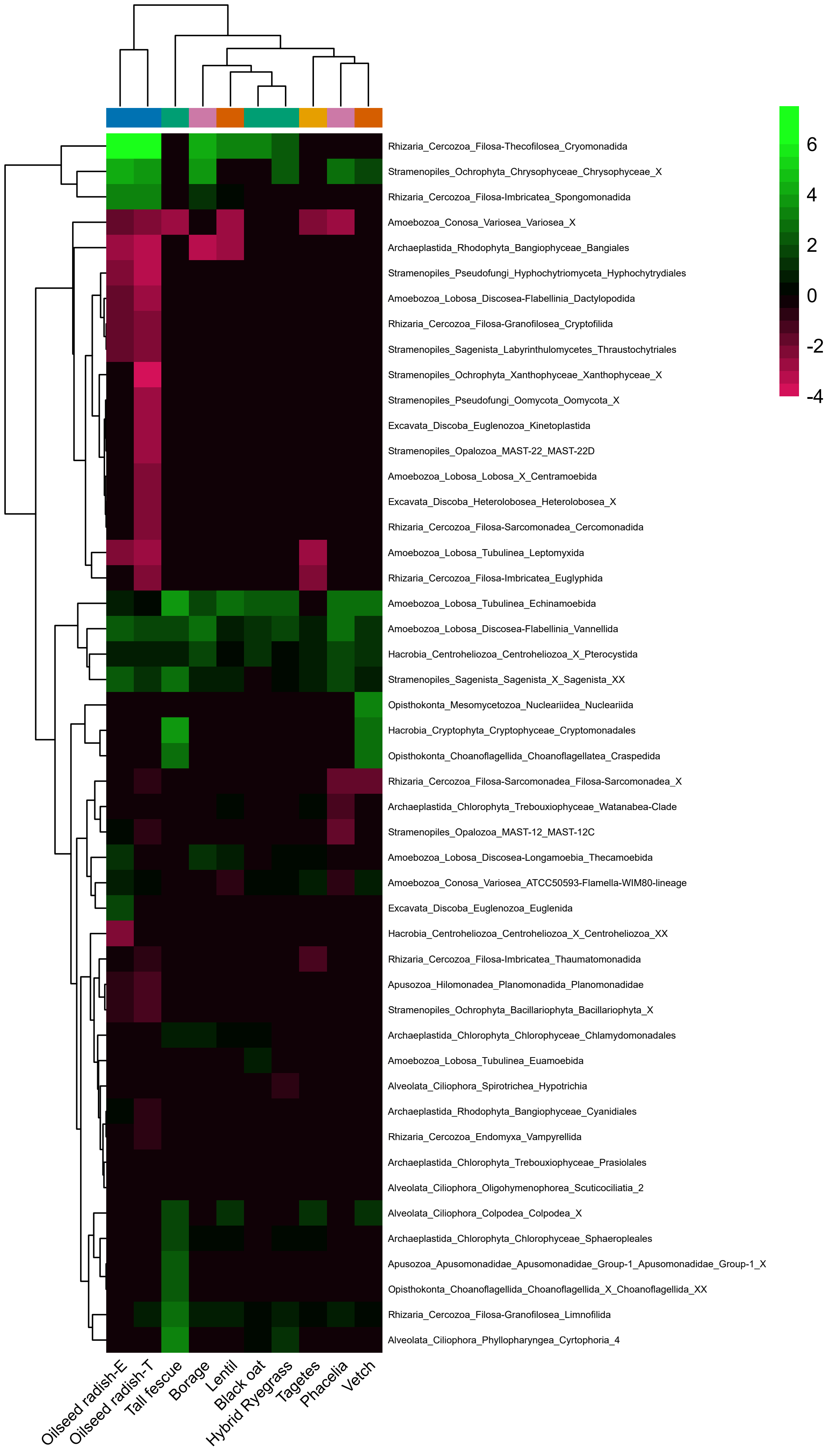
